# Cross-Tissue Transcriptomic Analysis Leveraging Machine Learning Approaches Identifies New Biomarkers for Rheumatoid Arthritis

**DOI:** 10.1101/2020.07.24.220483

**Authors:** Dmitry Rychkov, Jessica Neely, Tomiko Oskotsky, Steven Yu, Noah Perlmutter, Joanne Nititham, Alex Carvidi, Melissa Krueger, Andrew Gross, Lindsey A Criswell, Judith Ashouri, Marina Sirota

## Abstract

**Background/Purpose:** There is an urgent need to identify effective biomarkers for early diagnosis of rheumatoid arthritis (RA) and to accurately monitor disease activity. Here we define an RA meta-profile using publicly available cross-tissue gene expression data and apply machine learning to identify putative biomarkers, which we further validate on independent datasets.

**Methods:** We carried out a comprehensive search for publicly available microarray gene expression data in the NCBI Gene Expression Omnibus database for whole blood and synovial tissues from RA patients and healthy controls. The raw data from 13 synovium datasets with 284 samples and 14 blood datasets with 1,885 samples were downloaded and processed. The datasets for each tissue were merged, batch corrected and split into training and test sets. We then developed and applied a robust feature selection pipeline to identify genes dysregulated in both tissues and highly associated with RA. From the training data, we identified a set of overlapping differentially expressed genes following the condition of co-directionality. The classification performance of each gene in the resulting set was evaluated on the testing sets using the area under a receiver operating characteristic curve. Five independent datasets were used to validate and threshold the feature selected (FS) genes. Finally, we defined the RA Score, composed of the geometric mean of the selected RA Score Panel genes, and demonstrated its clinical utility.

**Results:** This feature selection pipeline resulted in a set of 25 upregulated and 28 downregulated genes. To assess the robustness of these FS genes, we trained a Random Forest machine learning model with this set of 53 genes and then with the set of 33 overlapping genes differentially expressed in both tissues and tested on the validation cohorts. The model with FS genes outperformed the model with common DE genes with AUC 0.89 ± 0.04 vs 0.87 ± 0.04. The FS genes were further validated on the 5 independent datasets resulting in 10 upregulated genes, *TNFAIP6*, *S100A8*, *TNFSF10*, *DRAM1*, *LY96*, *QPCT*, *KYNU*, *ENTPD1*, *CLIC1*, and *ATP6V0E1*, which are involved in innate immune system pathways, including neutrophil degranulation and apoptosis. There were also three downregulated genes, *HSP90AB1*, *NCL*, and *CIRBP*, that are involved in metabolic processes and T-cell receptor regulation of apoptosis.

To investigate the clinical utility of the 13 validated genes, the RA Score was developed and found to be highly correlated with the disease activity score based on the 28 examined joints (DAS28) (r = 0.33 ± 0.03, p = 7e-9) and able to distinguish osteoarthritis (OA) from RA samples (OR 0.57, 95% CI [0.34, 0.80], p = 8e-10). Moreover, the RA Score was not significantly different for rheumatoid factor (RF) positive and RF-negative RA sub-phenotypes (p = 0.9) and also distinguished polyarticular juvenile idiopathic arthritis (polyJIA) from healthy individuals in 10 independent pediatric cohorts (OR 1.15, 95% CI [1.01, 1.3], p = 2e-4) suggesting the generalizability of this score in clinical applications. The RA Score was also able to monitor the treatment effect among RA patients (t-test of treated vs untreated, p = 2e-4). Finally, we performed immunoblotting analysis of 6 proteins in unstimulated PBMC lysates from an independent cohort of 8 newly diagnosed RA patients and 7 healthy controls, where two proteins, *TNFAIP6/TSG6* and *HSP90AB1/HSP90*, were validated and the S100A8 protein showed near significant up-regulation.

**Conclusion:** The RA Score, consisting of 13 putative biomarkers identified through a robust feature selection procedure on public data and validated using multiple independent data sets, could be useful in the diagnosis and treatment monitoring of RA.

## Introduction

Rheumatoid arthritis (RA) is a systemic inflammatory condition characterized by a symmetric and destructive distal polyarthritis. Undiagnosed and untreated, RA can progress to severe joint damage, involve other organ systems, and predispose individuals to cardiovascular disease (Battisha et al., 2020; Lai et al., 2020). While our understanding of disease pathogenesis has greatly improved, and the number of available, effective therapeutics has significantly increased, there remains significant barriers to caring for patients with RA, and they continue to suffer from the morbidity and mortality associated with the disease. There is an urgent need to develop objective biomarkers for the early diagnosis and prompt initiation of disease-modifying therapy during the so-called “window of opportunity” (Bergstra et al., 2020; Burgers et al., 2019; Coffey et al., 2019; Van Nies et al., 2015). Additionally, clinicians need tests to help accurately assess disease activity or treatment targets in order to adjust therapy appropriately. Identification of biomarkers would greatly add to clinicians’ existing toolset used to evaluate patients with RA, helping to improve outcomes and alleviate the suffering caused by this prevalent disease.

Over the past decade, advances in genomic sequencing technology have greatly contributed to our understanding of inflammatory diseases and informed development of effective therapeutics. Transcriptomics provides a lens into the specific genes over- or under-expressed in a disease yielding insights into cellular responses. Given the numerous transcriptomic datasets that have been generated and made publicly available, there are now opportunities to combine these datasets in a meta-analytic fashion for unbiased computational biomarker discovery. Meta-analysis is a systematic approach to combine and integrate cohorts to study a disease condition which provides enhanced statistical power due to a higher number of samples when combined. Additionally, it provides an opportunity for leveraging all the disease heterogeneity combined from multiple smaller studies across diverse populations creating a more robust signature and better recognition of direct disease drivers as well as disease subtyping and patient stratification. Moreover, integrating datasets generated from the multiple target tissues within a given disease further strengthens the associations identified (Barbeira et al., 2019; Gomez-Cabrero et al., 2014; Haynes et al., 2020; Li et al., 2019; Lofgren et al., 2016; Pineda et al., 2020; L. Wang et al., 2015). This approach has been successfully applied to the study of antineutrophil cytoplasmic antibody (ANCA)-associated vasculitis (Friedman et al., 2019), dermatomyositis (Neely et al., 2019) and systemic lupus erythematosus (Haynes et al., 2020). These large datasets also present an opportunity to apply advanced machine learning techniques that were not previously feasible computationally, allowing for interrogation of the data with new and unbiased approaches.

Multiple studies have attempted to identify RA transcriptomic signatures in blood (Batliwalla et al., 2005; Sumitomo et al., 2018; Teixeira et al., 2009; L. Wang et al., 2015) and in synovial tissue (Asif Amin et al., 2017; Orange et al., 2018) separately or in cross-tissue analysis (Afroz et al., 2017; Li et al., 2014). The tissue-specific studies have found very few overlapping signals. The integrative meta-analysis studies combined a few datasets from each tissue (Afroz et al., 2017; Li et al., 2014) to identify an overlap of dysregulated genes and to recognize similarities and differences in disease pathways in both tissues. While this type of approach allows better understanding of the disease, a corresponding set of biomarkers is often redundant and requires extensive prioritization analysis and validation. Thus, more rigorous approaches for biomarker search with a built-in prioritization procedure are needed.

In this study, we leveraged publicly available transcriptomic datasets generated from microarray and RNA sequencing (RNA-seq) platforms from over 2,000 samples from whole blood and synovial tissue of patients with RA. After combining these datasets using a well-described meta-analytic pipeline (Hughey and Butte, 2015) and describing the expression pathways and cell types present in RA tissues, we developed and applied a robust machine learning and feature selection approach to identify unique and independent biomarkers which were subsequently refined and validated on test data. We then evaluated the diagnostic utility of this set of biomarkers and the correlation with disease activity measures to inform future clinical studies. The development of an effective blood test for the diagnosis and monitoring of RA can add valuable information to the physician’s assessment and help inform decision-making to improve the morbidity and quality of life for patients with RA.

## Methods

### Discovery data collection and processing

We carried out a comprehensive search for publicly available microarray data in the NCBI Gene Expression Omnibus (Barrett et al., 2013) (GEO) database (http://www.ncbi.nlm.nih.gov/geo/) for whole blood and synovial tissues in rheumatoid arthritis and healthy controls using the keywords “rheumatoid arthritis”, “synovium”, “synovial”, “biopsy”, and “whole blood”, among organisms “Homo Sapiens” and study type “Expression profiling by array” (**Figure 1a**). Datasets were excluded when samples were poorly annotated or run-on platforms with small numbers of probes. This search yielded 13 synovial datasets, which included 257 biopsy samples from subjects with RA and 27 from healthy controls obtained during joint or trauma surgeries (**Supplementary Table 1**). We identified 14 whole blood datasets with 1,885 samples: 1,470 RA patients and 415 healthy controls (**Supplementary Table 1**).

**Figure 1.**
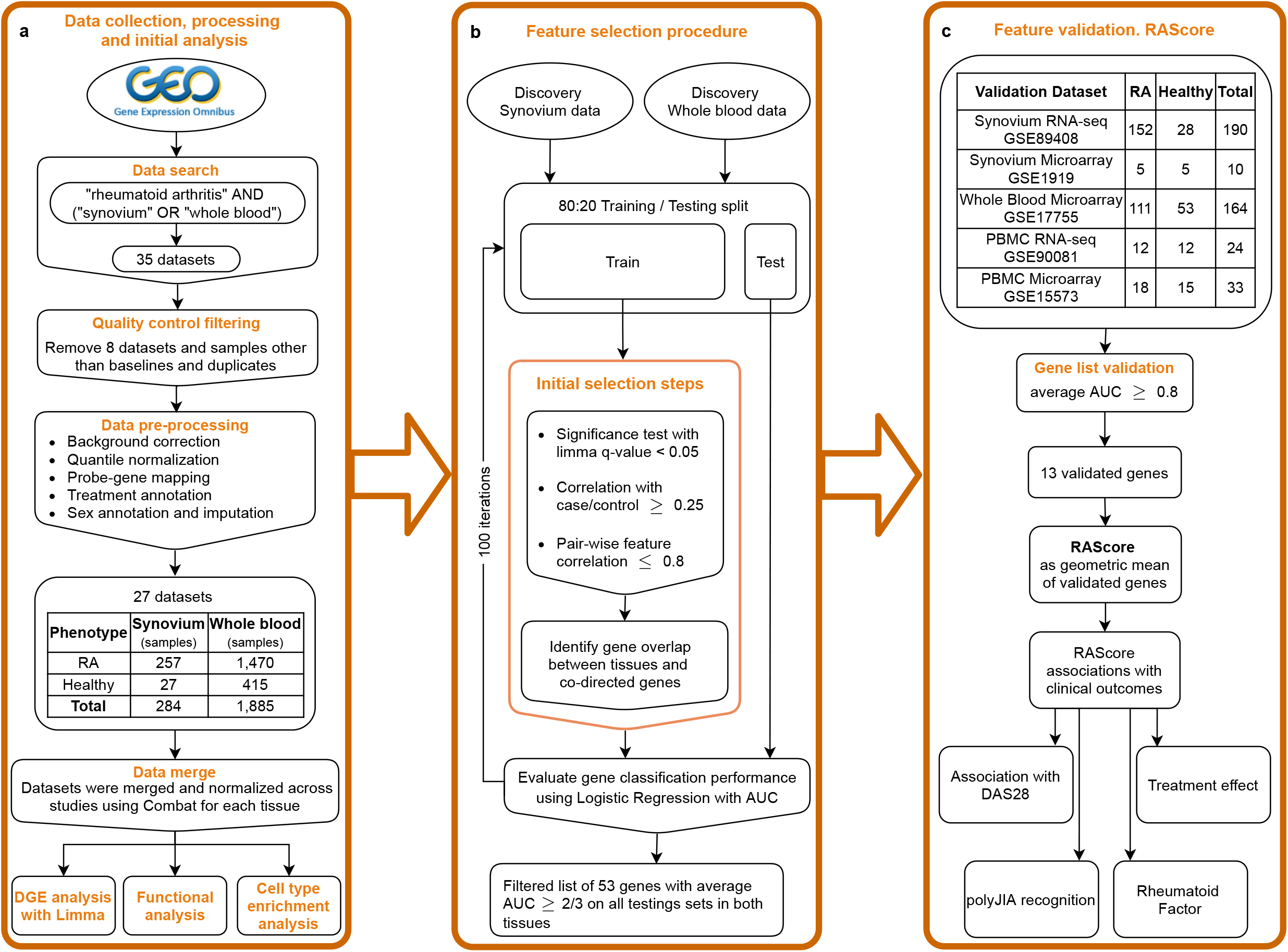
Study overview. a) Public data collection, processing and DGE analysis. b) Feature selection pipeline. c) Gene list validation on the independent datasets. Introducing the RAScore as a geometric mean of validated genes and its association with clinical outcomes.

Raw data was downloaded and processed using R language version 3.6.5 (R core team, 2018) and the Bioconductor (Huber et al., 2015) packages *SCAN.UPC* (Piccolo et al., 2012), *affy* (Gautier et al., 2004) and *limma* (Ritchie et al., 2015). Processing steps included background correction, log2-transformation, and intra-study quantile normalization (**Figure 1a**). Next, we performed probe-gene mapping, data merging and normalization across batches with Combat within the R package sva (Leek et al., 2012). The dimensionality reduction plots before and after normalization are shown in **Supplementary Figure 1**. After merging studies, the total number of common genes was 11,057 in synovium and 14,596 in whole blood.

### Validation data collection and processing

Five additional datasets from GEO were identified and downloaded: synovium microarray and RNA-seq, PBMC microarray and RNA-seq, and whole blood microarray datasets (**Supplementary Table 1**). Microarray data was processed as described above. RNA-seq data from GSE89408 were downloaded in a form of processed data of feature counts, which were normalized using the variance stabilizing transformation function *vst()* from the R package *DESeq2* (Love et al., 2014). RNA-seq data from GSE90081 were downloaded in a processed form of Fragments Per Kilobase Million (FPKM) counts, which were converted to Transcripts Per Kilobase Million (TPM) counts followed by log2 transformation with 0.1 offset.

### Differential gene expression and pathway analysis

Differentially expressed genes were identified using a linear model from the R package *limma* (Ritchie et al., 2015). To account for factors related to gene expression, the imputed sex and treatment categories were used as covariates. Treatment types were categorized based on the drug class (**Supplementary Table 2**). For 877 (40%) samples without sex annotations, sex was imputed using the average expression of Y chromosome genes. Significance for differential expression was defined using the cutoff of FDR p-value < 0.05 and abs(FC) > 1.2. Pathway analysis of differentially expressed genes was performed using the package *clusterProfiler* (Yu et al., 2012) with the Reactome database. To assess statistical significance of gene overlaps we computed p-values using the hypergeometric test.

### Cell type enrichment analysis

In order to estimate the presence of certain cell types in a tissue, we leveraged the cell type enrichment analysis tool, *xCell* (Aran et al., 2017) which computes enrichment scores for 64 immune and stromal cells based on gene expression data. We limited our analysis to 53 types of stromal, hematopoietic, and immune cells we expected to be present in blood and synovium. The cell types with a detection p-value greater than 0.2 taken as a median across all samples in a tissue were filtered. Non-parametric Wilcoxon-Mann test with multiple testing correction with Benjamini-Hochberg approach (cut-off 0.05) was used to assess significantly enriched cell types in synovium and whole blood in RA compared to healthy control subjects. The effect size of each cell type was estimated by computing the ratio of the mean enrichment score in RA patients over the mean score in healthy individuals.

### Feature selection pipeline

The feature selection procedure was partially described by Perez-Riverol, et al. (Perez-Riverol et al., 2017) and is represented in **Figure 1b**. First, for each tissue, the data were split into training and testing sets in an 80:20 ratio with random sample selection and class distribution preservation using the function *createDataPartition()* from the R package caret (Kuhn, 2008). Within each training set, a set of significant genes was identified using *limma* FDR p-value < 0.05. Pearson correlation coefficient was computed with the case-control status for each significant gene and those with r < 0.25 were filtered out. For robustness and reducing gene redundancy, we computed gene pair-wise correlations and removed genes with correlation greater than 0.8. Next, we overlapped the gene sets from both tissues and filtered out any genes differentially expressed in opposite directions in synovium and blood. To monitor statistical significance of gene overlaps we computed p-values using the hypergeometric test. To evaluate each gene’s performance in distinguishing RA from healthy samples, we trained a logistic regression model per gene on a training set for each tissue and tested on a testing set using area under receiver operating characteristic (AUROC) curve as a performance measure.

We repeated these steps 100 times to minimize bias of a random split into training and testing sets. From the resulting 100 gene sets, any gene that was found in each set in both tissues was further assessed. The AUC performance of each gene was averaged, and its standard deviation was calculated. We then set the AUC threshold to 2/3 and applied this criterion to the testing results to identify the genes with the best performance, the feature selected (FS) genes.

### Feature validation and RA Score

To evaluate and confirm superiority of the set of the FS genes over the set of the common DE genes, we trained machine learning models on the discovery blood data with these two gene sets and tested them on 5 independent data sets. As some genes were not present in all sets, the gene sets were reduced to the genes that were found in all 5 sets. We used three machine learning models: Logistic Regression, Elastic Net, and Random Forest, to compare the gene sets performance using AUROC.

Next, to further validate the FS genes identified in our pipeline, we trained a Logistic Regression model for each FS gene individually on the discovery blood data and tested on the validation sets (**Figure 1c**). AUROC was used as a performance measure. The genes with average AUC greater than 0.8 were selected. The selected genes were used to create the RA Score, computed by subtracting the geometric mean expression of the down-regulated genes from the geometric mean expression of the up-regulated genes.

Next, to assess the clinical value of the selected genes and the RA Score, we identified datasets with samples that included values for the disease activity score based on the 28 examined joints (DAS28) (Prevoo et al., 1995). We computed the Pearson correlation coefficients of the RA Score and expression levels of the RA Score genes with DAS28. Eight datasets with both RA and Osteoarthritis (OA) samples (**Supplementary Table 1**) were used to evaluate the ability of the RA Score to distinguish RA from OA. To report the summarized statistics as a combined p-value, Fisher’s method implemented in the R package *metap* was applied. The summarized odds ratio was computed by bootstrapping. GSE74143 was used to test the difference in RA Score between RA sub-phenotypes with and without rheumatoid factor by applying Student’s t-test. GSE45876 and GSE93272 were used to test the RA Score difference between treated and untreated RA patients via Student’s t-test. Additionally, we leveraged 10 datasets to test the ability of the RA Score to recognize polyarticular juvenile idiopathic arthritis (polyJIA), an inflammatory arthritis similar to RA affecting children under 16 years of age, by calculating the Odds Ratio (**Supplementary Table 1**).

### Immunoblot analysis

Patients with RA were recruited at University of California San Francisco Rheumatology Clinics. Blood samples and clinical measurements were obtained at the time of enrollment. Clinical measurements included Clinical Disease Activity Index (CDAI), erosion status (presence vs. absence), rheumatoid factor (positive vs. negative) and anti-cyclic citrullinated peptide (anti-CCP) autoantibodies (positive vs. negative). Healthy controls were recruited through local advertising and ResearchMatch (Harris et al., 2012), a national health volunteer registry that was created by several academic institutions and supported by the U.S. National Institutes of Health as part of the Clinical Translational Science Award (CTSA) program. Controls were matched to RA patients by age, gender, and race. Written informed consent was obtained from all participants and Institutional Review Board approval was obtained.

Frozen human PBMCs from 8 patients with RA (87.5% were seropositive) and 7 healthy controls were thawed, washed, resuspended in RPMI media. Those determined to have viability >89% (ViCell Counter) were used to make lysates for western blot. Cells were pelleted and then lysed by directly adding 10 % NP-40 lysis buffer to the final concentration of 1% NP40 (containing inhibitors of 2 mM NaVO4, 10 mM NaF, 5 mM EDTA, 2 mM PMSF, 10 μg/ml Aprotinin, 1 μg/ml Pepstatin and 1 μg/ml Leupeptin) as previously described (Ashouri et al., 2019). Lysates were placed on ice and centrifuged at 13,000 g to pellet cell debris. Supernatants were mixed with a 6X loading buffer containing BME. Proteins were separated on 10% Bis-Tris gels (Thermo Fisher) and transferred to Immobilon-P polyvinylidene difluoride membranes (Millipore) via standard tank transfer techniques. Primary staining was performed with the following antibodies: TSG-6 (Santa Cruz Biotechnology: sc-377277, clone: E-1), Protein S100A8/Calgranulin A (Santa Cruz Biotechnology: sc-48352, clone: C-10), CD39 (MyBiosource: MBS2541905), HSP90beta (Cell Signaling Technology: #5087), Ly96/MD-2 (Novus Biologicals: NB100-56655), TRAIL (Cell Signaling Technology: #3219S). Membranes were blocked using a TBS-T buffer containing 2% BSA, and probed with primary antibodies as described, overnight at 4 °C. The following day, blots were rinsed and incubated with HRP-conjugated secondary antibodies. Horseradish peroxidase (HRP)-conjugated secondary antibodies from Southern Biotech and blots were visualized with SuperSignal ECL reagent or SuperSignal West Femto maximum sensitivity substrate (Pierce Biotechnology) on Chemi-Doc Image Lab station (Bio-Rad).

Each protein was measured on a set of two immunoblots and normalized to the beta-actin level. In order to combine and normalize measured protein amounts from both blots for further analysis, we applied an empirical Bayes approach *ComBat* implemented in R package *sva* (Leek et al., 2012). Each of two pairs of control replicates were averaged. The Mann–Whitney–Wilcoxon test was used to compare groups and unadjusted p-values were reported.

## Results

### Cross-Tissue Differential Expression and Pathway Analysis Reveals Significant Similarities on Gene and Pathway Levels

The differential gene expression analysis identified 1,389 genes with 789 up-and 600 down-regulated genes in the synovium (**Supplementary Figure 2ab** and **Supplementary Table 3**) and 155 genes with 110 up- regulated and 45 down-regulated genes in the blood (**Supplementary Figure 3ab** and **Supplementary Table 4**). The pathway analysis revealed that in both tissues, up-regulated genes shared enrichments in innate immune system, neutrophil degranulation, interferon signaling, cytokine signaling, toll-like receptor (TLR) cascades, regulation of TLR by endogenous ligand, and caspase activation via extrinsic apoptotic signaling pathways (**Figure 2a, Supplementary Figures 2cd, 3cd, Supplementary Tables 5,6**). However, interferon gamma signaling, immunoregulatory interactions between a lymphoid and non-lymphoid cell, PD-1 signaling were specific for synovium (**Supplementary Table 5**), whereas apoptosis, programmed cell death, antiviral mechanisms, caspase activation via death receptors in the presence of ligand were specific for blood (**Supplementary Table 6**). The down-regulated genes were commonly involved only in the interleukin-4 and interleukin-13 signaling pathways (**Figure 2b, Supplementary Figures 2ef, 3ef**). Some of the pathways were not shared, suggesting the existence of distinct underlying molecular mechanisms operating in tissues. For example, signaling by interleukins, TCR signaling, and MHC class II antigen presentation pathways were specific only for synovium (**Supplementary Tables 5,6**). The latter was fully consistent with our previous work demonstrating enrichment of Nur77 – a specific marker of TCR signaling – in joint infiltrating CD4+ T-cells, suggesting that CD4+ T-cells are recognizing intra-articular antigen (Ashouri et al., 2019).

**Figure 2.**
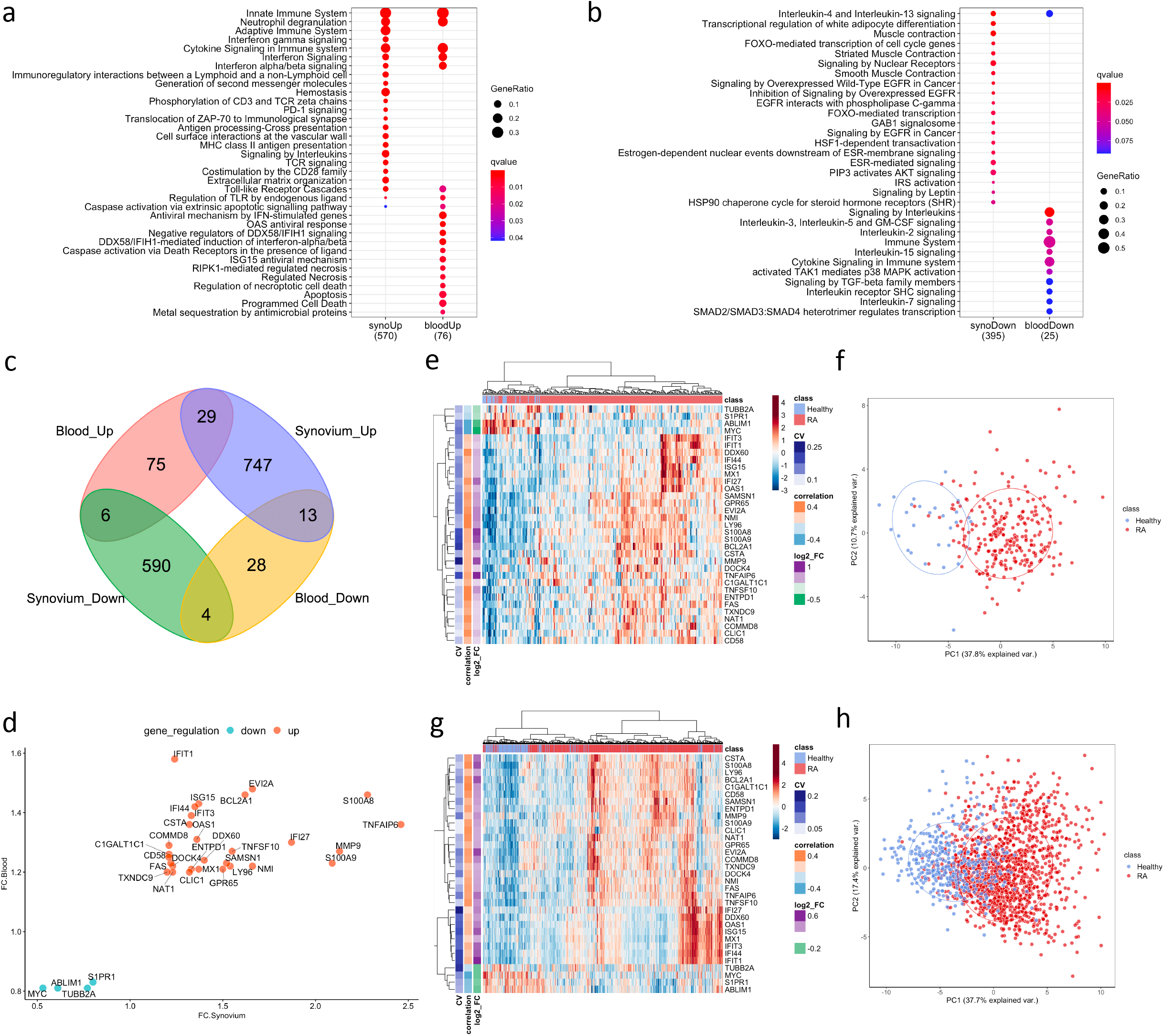
DE genes overlapped between synovium and whole blood tissues. Top Reactome common and different pathways for a) Up-regulated and b) down-regulated genes. Venn diagram of up- and down-regulated genes in synovium and blood: 29 common up-regulated genes (p = 3e-09) and 4 common down-regulated genes (p = 0.28). d) Comparison scatter plot of fold changes between common genes in synovium and blood. Heatmap and PCA plots of common genes in e) and f synovium and f) blood. Vertical bars in the heatmap plots represent the color-coded coefficients of variation, Pearson correlations and log2 fold changes.

When evaluating the overlap between differentially expressed genes in synovium and blood, there were 29 genes commonly up-regulated: *TNFAIP6, S100A8, MMP9, S100A9, IFI27, EVI2A, NMI, BCL2A1, TNFSF10, LY96, SAMSN1, GPR65, DDX60, ISG15, MX1, OAS1, IFI44, ENTPD1, IFIT3, CSTA, CLIC1, IFIT1, DOCK4, NAT1, FAS, C1GALT1C1, CD58, COMMD8, TXNDC9*; and 4 down-regulated genes: *S1PR1, TUBB2A, ABLIM1*, and *MYC* (**Figure 2c, Supplementary Table 7**, hypergeometric test p = 3e-9). However, the overlap of down-regulated genes did not meet statistical significance (hypergeometric test p = 0.28, **Figure 2d**). The common differentially expressed (DE) genes formed more distinct clusters of RA and control samples for both synovium (**Figure 2ef**) and blood (**Figure 2gh**) than all DE genes for these tissues (Supplementary **Figure 2ab** and **Supplementary Figure 3ab**). The enriched Reactome pathways of these common up-regulated genes included interferon signaling, neutrophil degranulation, regulation of TLR by endogenous ligand and caspase activation via extrinsic apoptotic signaling pathway and via death receptors in the presence of ligand, whereas down-regulated genes are associated with Interleukin-4 and 13 signaling and cell cycle pathways. These results were consistent with the pathway analysis above.

### Cell-Type Deconvolution Analysis Identifies a Reverse Signal in Blood and Synovium

The cell type enrichment analysis with xCell in synovium revealed the enrichment of immune cell types, including, CD4+ and CD8+ T-cells, B-cells, macrophages and dendritic cells in RA samples (**Figure 3a**). However, opposite but weaker associations were seen in whole blood samples with enrichment of T- and B-cells in healthy controls (**Figure 3b**). Lymphocytes, including CD8+ T cells and B cells, were significantly enriched in both tissues, however, these were enriched in opposite directions with enrichment in cases in synovium but enrichment in controls in blood (**Figure 3c**). This finding was confirmed in validation datasets (**Figure 3d**). The significant cell types in synovium and blood showed high correlations in validation data: r = 0.71 (p = 1.3e-5) for synovium (**Figure 3e**) and r = 0.61 (p = 0.004) in blood (**Figure 3f**).

**Figure 3.**
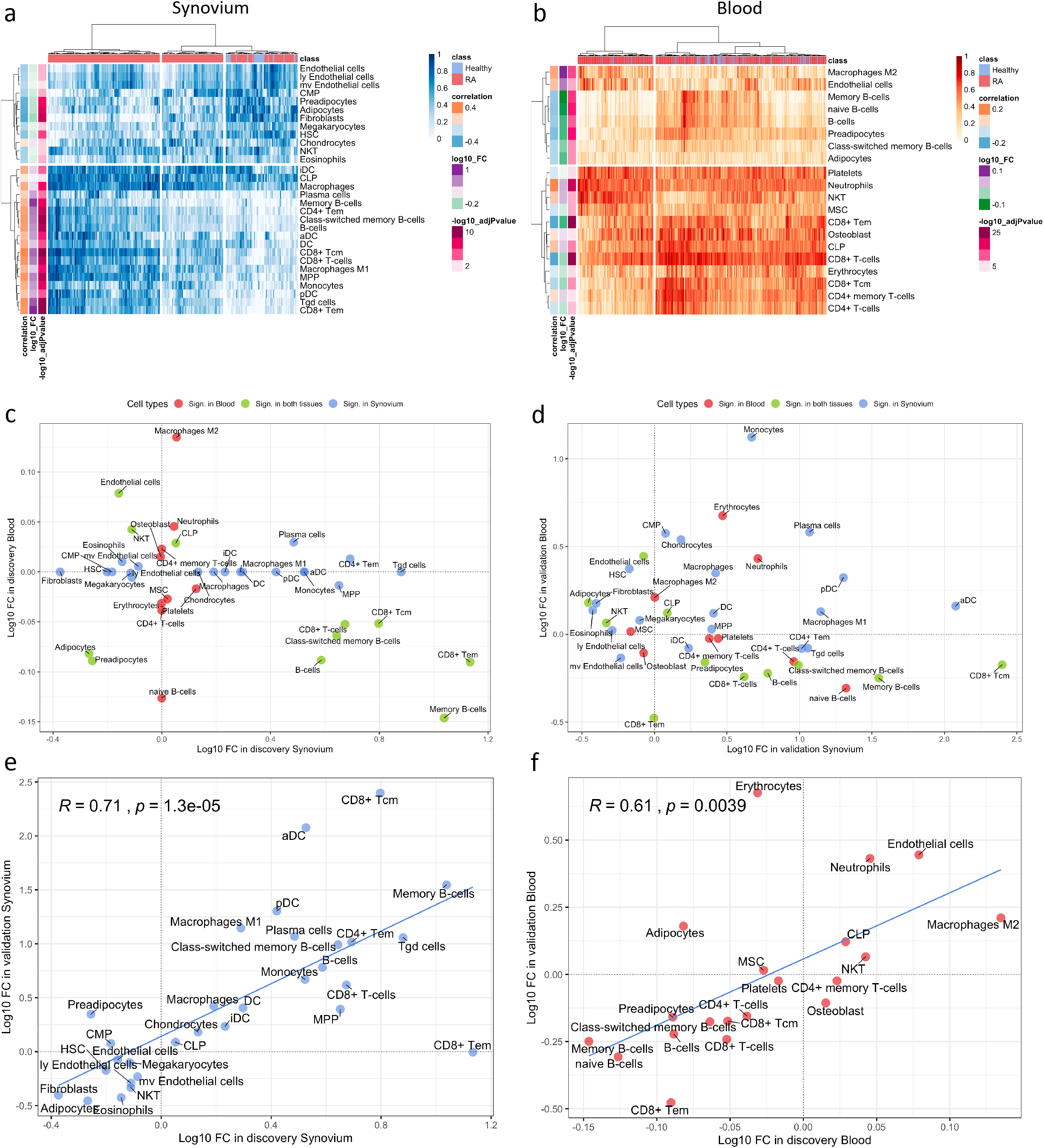
Cell type enrichment analysis for synovium and and whole blood tissues. BH adj p-values <0.05. 30 significant cell types in synovium, 20 significant cell types in WB, 11 common significant cell types.

### Machine Learning Feature Selection Strategy to Identify Robust Cross-Tissue Biomarkers of RA

Aiming to determine a more robust list of putative biomarkers that are strongly associated with RA in both synovium and whole blood tissues and have higher predictive power, we applied an iterative feature selection procedure leveraging the gene expression data from both tissues. In the pipeline, only 10,071 genes that were common between synovium and whole blood data were used. At each iteration, only genes found significantly dysregulated in both tissues following the condition of co-directionality were kept (p = 6.3e-10). As a result of these filtering steps, on average 65 up-regulated and 71 down-regulated were selected from each iteration (See **Methods**). From 100 iterations, any gene significantly dysregulated in all the iterations was selected, resulting in a set of 53 genes: 25 upregulated and 28 down-regulated (**Supplementary Table 8**). A summary of the average AUC performance from the 100 iterations for each gene are shown in the **Figure 4a** and **Supplementary Table 8**. The AUC for selected genes in synovial tissue varied with mean 0.853 ± 0.005 for training and 0.866 ± 0.006 for testing sets, whereas for the whole blood the mean AUC was 0.744 ± 0.006 for both training and testing sets.

**Figure 4.**
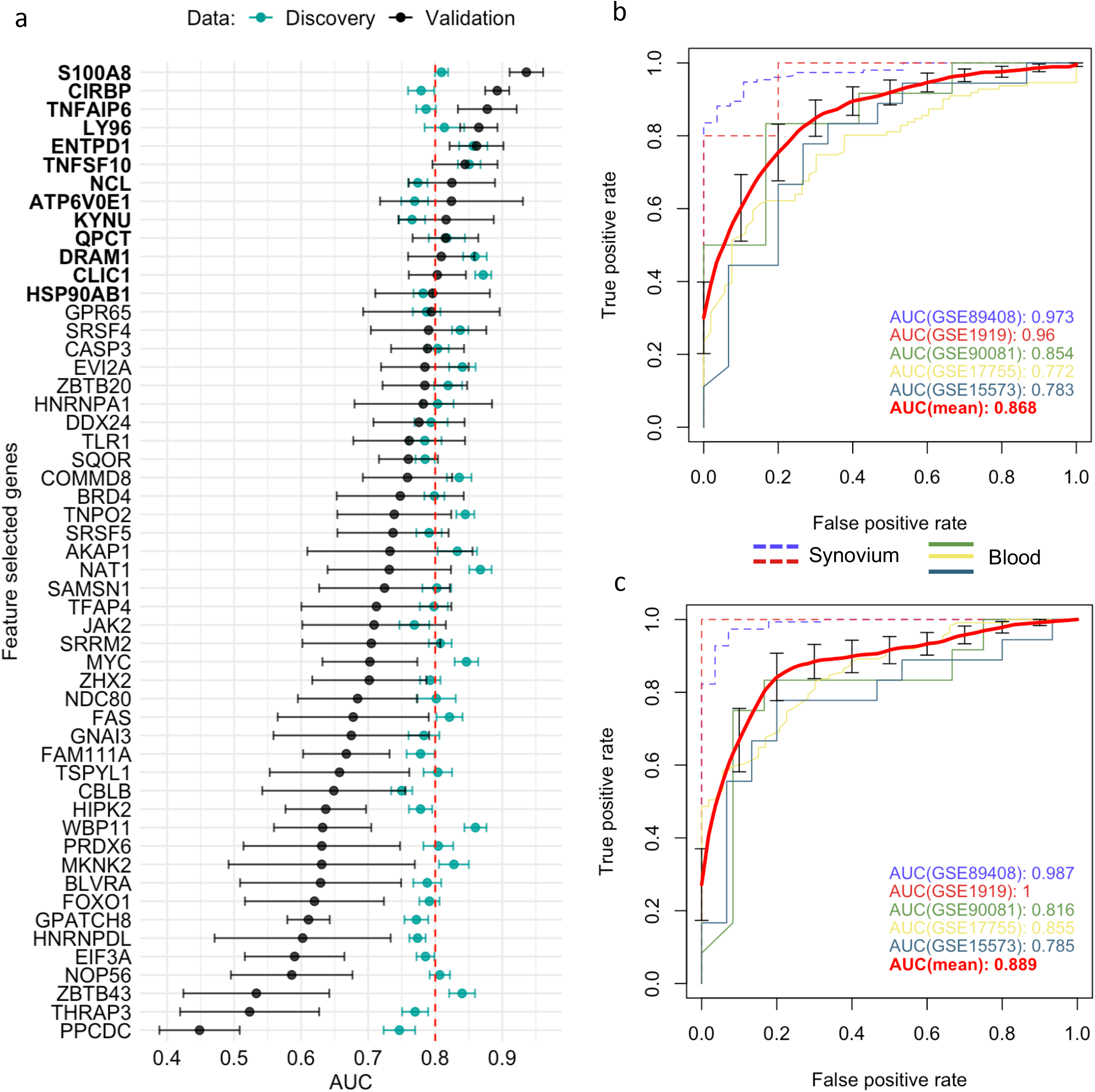
Feature selected genes. a) Mean AUC performance with standard error for each feature selected gene on testing synovium and blood datasets (green) and on five independent validation datasets (black). 13 genes with AUC greater than 0.8 on validation datasets were chosen as the best performing genes. Mean AUC performance with standard errors of a RF model trained on discovery blood data with b) common DE genes and c) feature selected genes on five independent validation datasets.

We leveraged 5 publicly available independent datasets on synovium and blood to validate these results (see Methods) (**Supplementary Table 1**). First, we compared the classification performance of the set of 53 feature selected genes to the set of 33 common DE genes (**Methods**). Since not all genes were measured across the validation studies, the sets were reduced to 26 of 33 common DE genes and 38 of 53 feature selected genes. We found the set of 38 feature selected genes has superior performance over the set of 26 common DE genes for all three ML methods (Supplementary Figure 4). The largest difference in performance was for the Random Forest model: the model with the common DE genes had an AUC of 0.868 ± 0.043 (95% CI [0.785, 0.951]) (**Figure 4b**), while the model with the feature selected genes performed with 0.889 ± 0.044 (95% CI [0.811, 0.966]) (**Figure 4c**).

Next, the set of 53 feature selected genes was thresholded to an average AUC of 0.8 using the validation sets resulting in 10 up-regulated *TNFAIP6, S100A8, TNFSF10, DRAM1, LY96, QPCT, KYNU, ENTPD1, CLIC1*, and *ATP6V0E1*, and three down-regulated *HSP90AB1, NCL*, and *CIRBP* genes (**Figure 4a, Supplementary Figure 5**). Five of the related proteins, *TSG-6, MRP-8/Calgranulin-A, TNFSF10/TRAIL, Ly-96*, and *QC*, are known to be normally secreted into blood, while Kynureninase, *HSP 90-beta* and *CLIC1* are localized to cytosol (Uhlén et al., 2019).

### Clinical Implications of Transcription Based Disease Score

In order to assess the clinical utility of the 13 validated genes, we introduced a scoring function, RA Score, which is derived by subtracting the geometric mean of expression values of down-regulated genes from the geometric mean of up-regulated genes. With this definition, the RA Score is 2-fold (95% CI [1.8, 2.2], p = 3e-15) larger for RA in comparison to healthy samples in synovium. In whole blood, the RA Score has a mean effect size of 1.37 (95% CI [1.34, 1.4], p = 1e-108). In validation datasets, the RA Score had a mean effect size of 5.5 in synovium (95% CI [3.8, 8.2], p = 1e-10) and 2.4 in blood (95% CI [2.1, 2.8], p = 3e-23). This score showed utility in monitoring disease activity, diagnostics, and treatment response and also was generalizable to both RF-positive and RF-negative RA as well as polyJIA.

Four datasets with 411 samples included in our meta-analysis had available disease activity score (DAS28) annotations. Assessing the correlation with DAS28 for each gene individually, the most positively correlated gene was *S100A8* with mean R = 0.28 (95% CI [0.19, 0.37]) and most anti-correlated gene *HSP90AB1* with mean r = −0.23 (95% CI [−0.32, −0.14]) (**Figure 5b, Supplementary Figure 6**). The RA score performed better than any single gene, positively correlated with DAS28 where the average correlation was 0.33 with 95% CI [0.24, 0.41] (**Figure 5a**), suggesting this score could be helpful as a disease activity biomarker. We also determined the correlation of the RA Score with DAS28 in these datasets and obtained Pearson correlation coefficients from 0.25 to 0.43 in blood and 0.31 in synovium (**Supplementary Figure 7**).

**Figure 5.**
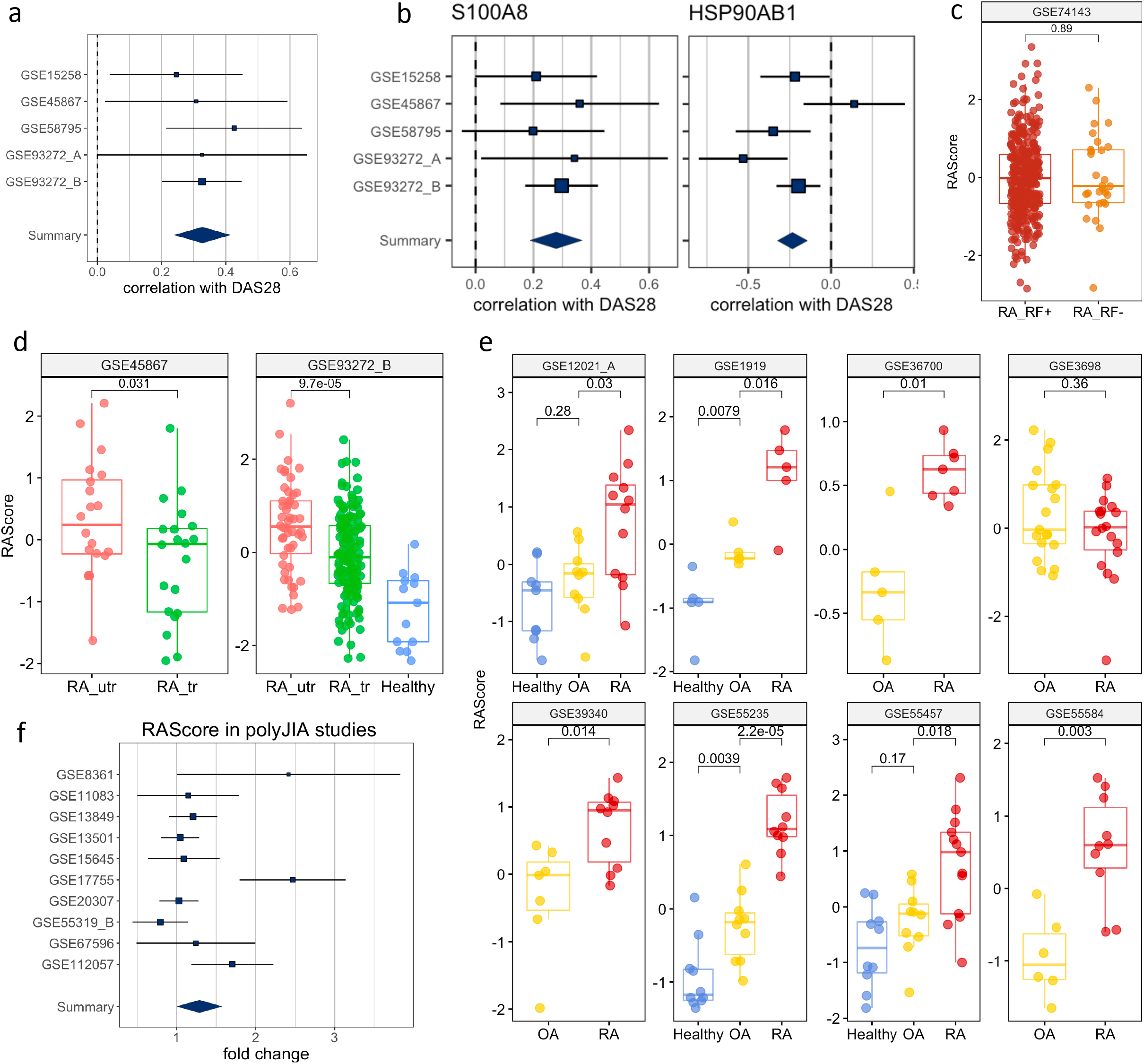
Clinical Interpretation of the RAScore. a) A forest plot of correlation RAScore with DAS28. b) Forest plots of correlations of some RAScore Panel genes with DAS28. c) RAScore tracks the treatment effect in both synovium and blood but shows no difference between RF− and RF-phenotypes. d) RAScore tracks the treatment effect in both synovium and blood. RA_utr: RA untreated. RA_tr: RA treated. e) RAScore distinguishes Healthy, OA and RA samples in synovium. f) RAScore distinguishes Healthy and polyJIA samples. The p-values were obtained using Student’s t-test.

To investigate the ability of the RA Score to differentiate RA from osteoarthritis (OA), we identified eight datasets that had both RA and OA samples available. **Figure 5e** shows the distributions of RA Score for RA, OA and healthy samples in eight available datasets. In most datasets, the RA Score was able to significantly differentiate OA from RA (OR 0.57, 95% CI [0.34, 0.80], p = 8e-10) and healthy samples (OR 1.53, 95%CI [1.37, 1.69], p = 7.5e-4) suggesting that this score could be useful diagnostically.

One dataset in whole blood, GSE74143, had annotations for RF-positivity. The RA Score performed similarly in both RF-positive and RF-negative RA samples suggesting the applications of this score are generalizable to these RA subtypes (t-test, p = 0.9) (**Figure 5c**). Furthermore, we tested the utility of this score in 10 datasets from polyJIA given that this subtype of JIA is most similar to RA, and also found comparable performance in the ability to differentiate polyJIA from healthy controls (OR 1.15, 95% CI [1.01, 1.3], p = 2e-4) (**Figure 5f**). Thus, this score could also be useful in the pediatric arthritis population.

Lastly, it appears the RA Score also tracks with treatment response. In two datasets, RA patients had transcriptional measurements before and after treatment with disease-modifying antirheumatic drugs (DMARD). The RA score significantly (p = 2e-4) decreases between pre- and post-treatment measurements (**Figure 5d**).

### Western Blot Validation

We next examined whether we could validate differences in transcript expression by protein in patients with newly diagnosed RA (validation cohort) prior to treatment initiation based on our RA Score results. Six candidate proteins, for which commercial antibodies are available, were selected by highest absolute fold changes based on transcript expression visible in PBMC’s, as well as in synovial tissue, from our discovery data. Protein levels were assayed using mixed PBMC lysates from the validation cohort (**Figure 6, Supplementary Figure 8**). Notably, the TSG6 protein was significantly upregulated in RA PBMCs (**Figure 6b**), while HSP90 was significantly downregulated in the RA validation cohort compared to controls (**Figure 6d**), supporting findings from the transcriptional discovery data. Similar to our RA Score findings, S100A8 was upregulated in some RA samples, though it did not reach statistical significance. Notably, Ly96 protein levels trended in the opposite direction compared to the transcriptional dataset (**Figure 6b**).

**Figure 6.**
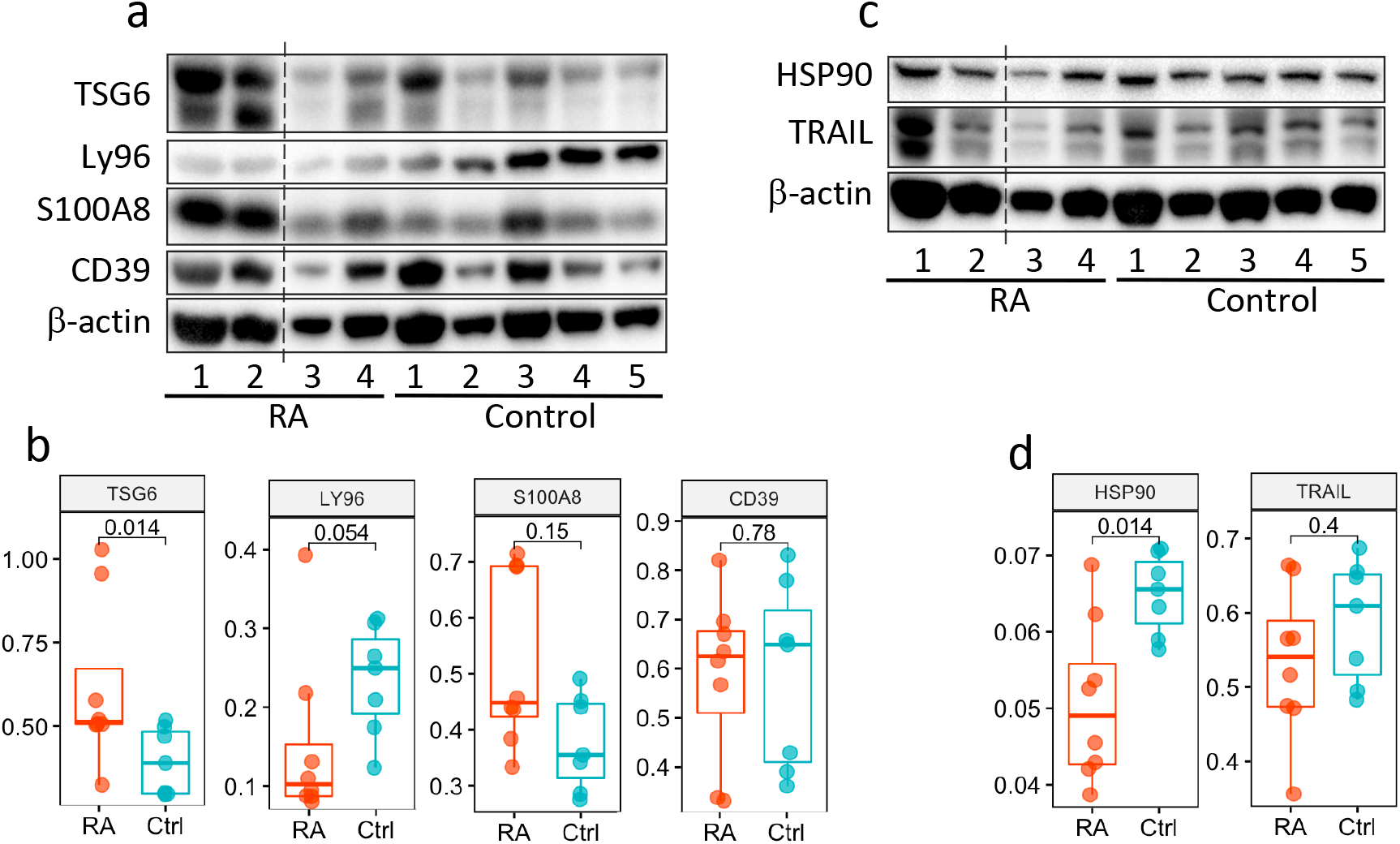
Validation of RAScore proteins. (**a** and **c**) Immunoblot analysis of 6 RAScore proteins in unstimulated PBMC lysates from subjects with RA (n=4) and healthy controls (n=5). Data representative of 2 immunoblots. (**b** and **d**) Box plots are quantification of RAScore protein levels normalized to GAPDH pooled from 2 immunoblot experiments (as shown in **a**, **c**, and Supplemental Fig 8); RA (n=8) and healthy control (n=7) samples. Significance determined by Mann-Whitney-Wilcoxon test, (**b** and **d**). Dotted line represents lanes removed from non-RA subjects, otherwise immunoblots **a** and **d** are montages of the same western blot.

## Discussion

In this study, we leveraged publicly available microarray gene expression data from both synovium and peripheral blood tissues in search of putative biomarkers for RA. The cell type enrichment analysis revealed the prevalence of lymphocytes in RA synovial tissue, in contrast to RA blood, likely due to tissue infiltration of immune cells as well as their homing to lymph organs (lymph nodes and spleen). This observation additionally supported our study objective of leveraging the data from both synovium and blood to identify genes expressed concurrently in both tissues. We first applied a conventional approach (Dudley et al., 2009; Hao et al., 2017; Shchetynsky et al., 2017; L. Wang et al., 2015; Wu et al., 2010) of intersecting the differentially expressed genes from both tissues and obtained a list of 33 common genes. Some of our results showed agreement with previous studies, identifying similar biological processes (e.g., cytokine signaling, immune and defense response, response to biotic stimulus) (Afroz et al., 2017; Teixeira et al., 2009; L. Wang et al., 2015). Furthermore, the differentially expressed genes common to both tissues better distinguished RA cases from healthy controls than all differentially expressed genes combined. While this list of overlapping genes provides valuable insight into disease biology, the predictive ability can be further improved by applying more advanced machine learning methods to prioritize candidate markers and remove redundancy.

Our specific machine learning method identified a robust and non-redundant set of biomarkers concurrently expressed in both RA target tissues. This resulted in 53 protein-coding genes that outperformed the set of the common DE genes in outcome prediction tasks using independent data. In further validation steps, we identified and selected 10 up- and three down-regulated genes with the highest performance. The up-regulated genes are highly expressed in diseased synovial tissue, and their elevated protein levels in blood may represent RA biomarkers. However, the combination of these 13 validated genes into a transcriptional gene score, the RA Score, performed better in clinical applications than any one gene and could potentially serve as a clinical blood test for accurate disease diagnosis and monitoring disease activity and response to treatment. Furthermore, this score performed similarly in RF-positive and RF-negative RA and also distinguished polyJIA from healthy controls, broadening the scope of possible clinical applications. Some genes/proteins, e.g. S100A8/MRP8, ENTPD1/CD39, KYNU, and TNFAIP6, from the score were previously found to be associated with JIA (Bojko, 2017; Brachat et al., 2017; Griffin et al., 2009; Hinze et al., 2019; Holzinger et al., 2019; Jiang et al., 2013; Korte-Bouws et al., 2019; Moncrieffe et al., 2010). Treatment effect was also captured with a significantly lower RA Score for DMARD-treated patients compared to untreated patients. Moreover, since the genes were identified from both blood and synovium and followed the condition of co-directionality, i.e., upregulated or downregulated in both tissues, the RA Score test based on blood only becomes tenable.

The 13 genes identified using these machine learning methods in the feature selection pipeline represent candidate biomarkers in RA. Six of the 13 RA Score Panel genes (TNFAIP6, S100A8, TNFSF10, LY96, ENTPD1, and CLIC1) were also among the 33 common DE genes, whereas seven of the 13 RA Score Panel genes (DRAM1, QPCT, KYNU, ATP6V0E1, NCL, CIRBP and HSP90AB1) were not. Many of these genes have been described in the literature and studied in the context of RA demonstrating biologic plausibility of the RA Score.

***TNFAIP6***, also known as TSG-6, encodes a secretory protein that is produced in response to inflammatory mediators, with high levels detected in the synovial fluid of patients with RA and OA (Wisniewski et al., 1993). TNFAIP6 is thought to play an anti-inflammatory role in arthritis and protect destruction of joint cartilage, which has been demonstrated in many arthritis mice models (Milner and Day, 2003). ***S100A8*** is a calcium binding protein that forms a heterodimer with S100A9 known as calprotectin (S100A8/A9). Calprotectin is constitutively expressed in neutrophils and monocytes but massively upregulated during inflammatory responses as an important mediator of inflammation (Wang et al., 2018). Thus, it has been extensively studied as a potential biomarker in several inflammatory diseases including RA. A study investigating the use of calprotectin as measured in sera of RA patients with moderate to severe disease found an association with disease activity, though it was less useful in monitoring radiographic disease progression or treatment response (Nordal et al., 2016). Of note, while the 33 common DE Genes set includes S100A9, this gene is not one of the 53 FS genes or subsequent 13 RA Score Panel genes likely because our process excludes genes that are highly correlated (pairwise feature correlation greater than 0.8). ***TNFSF10***, also known as TNF-Related Apoptosis Inducing Ligand (TRAIL), encodes a protein that induces apoptosis of tumor cells but is also of interest in RA as it has been thought that TNSF10 might induce apoptosis of hyperplastic synoviocytes and reduce immune cell hyperactivity thus providing a protective effect to the joint. However, this is still controversial as there is also evidence that TNSF10 may promote joint destruction and exacerbate RA (Audo et al., 2013). ***LY96***, also known as MD2, encodes a protein which often is a coreceptor with TLR4 forming the TLR4/MD2 complex (Nagai et al., 2002) and has previously been found to be upregulated in patients with rheumatoid arthritis (Brentano et al., 2005; Manček-Keber et al., 2015; Ospelt et al., 2008). ***ENTPD1***, also known as CD39, is a gene found to be an expression quantitative trait locus associated in RA affecting levels of CD39+, CD4+ regulatory T-cells (Spiliopoulou et al., 2019). In RA, low levels of CD39+ expressing Tregs were associated with methotrexate resistance suggesting this could be a biomarker to predict responders and non-responders (Gupta et al., 2018; Peres et al., 2015; Zacca et al., 2020).

Seven of the 13 RA Score Panel genes (DRAM1, QPCT, KYNU, ATP6V0E1, NCL, CIRBP and HSP90AB1) were uniquely identified by our machine learning pipeline and not identified by the traditional DE gene overlap method. Four of these genes, KYNU, QPCT, CIRBP, and HSP90AB1 have previously been associated with RA. ***KYNU*** encodes the Kynureninase enzyme which is involved in a pathway of tryptophan metabolism related to immunomodulation and inflammation (Harden et al., 2016; Q. Wang et al., 2015). KYNU expression has been found to be increased in chondrocytes and synovial tissue of RA patients compared to healthy patients (Chen et al., 2018). Moreover, increased tryptophan degradation has been observed in the blood of RA patients (Schroecksnadel et al., 2003). The gene ***QPCT*** encodes Glutaminyl-Peptide Cyclotransferase, or QC for short (NCBI, 2020a), which plays a role in maintaining inflammation (Kehlen et al., 2017). A clinical study found that expression of QPCT was significantly increased in the blood and in the fluid from gingival crevices of RA patients compared to healthy controls (Bender et al., 2019). ***CIRBP*** encodes cold-inducible RNA-binding protein which can be induced under stress and have a cytoprotective role but has also been increasingly recognized for participating in proinflammatory response (Liao et al., 2017). CIRBP binds to the TLR4/MD2 complex of macrophages and monocytes in the circulation or tissues, thereby activating the NF-Kappa B pathway and resulting in the release of inflammatory mediators (Liao et al., 2017; Qiang et al., 2013). A clinical study measured CIRBP mRNA expression of CD14+ monocytes of five healthy and five RA patients and found the relative expression of CIRBP was higher in RA patients (Yoo et al., 2018), whereas our analysis found down-regulated CIRBP expression in whole blood from RA compared to healthy control. Further study of CIRBP in RA patients, particularly single cell analysis, is warranted. ***HSP90AB1*** encodes the protein HSP90β (NCBI, 2020b). Post-translationally citrullinated isoforms of heat shock protein 90, including citHSP90β, have been identified as potential autoantigens in patients with RA-associated interstitial lung disease (Harlow et al., 2014; Travers et al., 2016). We did not find evidence of an association between DRAM1, ATP6VOE1, and NCL, however, these may represent novel genes that warrant future study. DRAM1 and NCL have been shown to be associated with other autoimmune diseases, such as Systemic Lupus Erythematosus (Molineros et al., 2013; Yang et al., 2013) and Multiple Sclerosis (Liu et al., 2012). Furthermore, DRAM1 is a gene involved in autophagy, an important mechanism for regulating the immune response and autophagy modulation has been postulated as a potential therapeutic in RA (Vomero et al., 2018).

We examined six proteins from RA Score using the immunoblotting technique and confirmed two of them, TNFAIP6/TSG6 and HSP90AB1/HSP90, with S100A8/MRP8 protein trending near significance. Our findings for these proteins highlight that the generation of the RA Score by which to help diagnose patients from RA could be helpful in clinical practice. Further validation studies are underway to examine the transcriptional profile of RA PBMCs on a single cell level, in which transcript changes can be assessed based on cell type, followed by protein validation studies in more RA subjects looking at lysates sorted from specific immune cell subtypes identified by single cell analysis. This will allow us to further fine tune the RA Score genes identified by our analysis. Indeed, studies examining the relationship between protein and mRNA levels highlight the complexity of gene expression regulation (Liu et al., 2016).

Several limitations of this study should be recognized. Several datasets, especially in whole blood, had significantly more cases than controls with some datasets containing no healthy controls. This significant class imbalance might result in class variance imbalance and lead to increased type 1 and type 2 errors. To at least partially address this limitation, we included two datasets of healthy individuals to enrich the blood data with control samples. Likewise, the validation cohorts also had an imbalance of cases and controls and two out of three were from PBMC rather than whole blood. The latter was also the case for the validation cohort in the immunoblotting analysis. This could possibly lead to lower AUC performance for genes in the validation datasets but likely does not overestimate the performance of our genes. Additionally, not all samples were annotated for important covariates such as sex and medication use. All sample annotations were kept from the original publications, though for 40% of samples the sex annotations were not available, and they were imputed based on the expression levels of Y chromosome genes. Likewise, most case samples were from RA patients who were taking various medications. Even though the treatments were used in the differential expression analysis as covariates (including untreated patients) there still exists the possibility of confounding.

In this study, we present a robust machine learning pipeline to search for putative biomarkers: each gene went individually through a feature selection procedure with multiple iterations on the discovery data and was independently tested on the validation cohorts. The gene redundancy was decreased selecting the best performing genes in RA association prediction. We apply the pipeline to a set of over 2000 samples to identify the RA Score as a potential diagnostic panel and show its clinical utility. The strength of RA Score is in the independence of its composite genes. Further development of the RA Score as a clinical tool requires greater understanding and validation of its composing genes with experimental analysis of the protein levels in RA patients and healthy individuals through prospective trials. An independent longitudinal study would bring better understanding of the early diagnostic and disease monitoring capability of the proposed panel. Additional experiments leveraging single cell technologies will further enhance our understanding of the biology and cell type specific effects in RA allowing us to refine the proposed diagnostic strategies.

## Acknowledgements

This work was in part supported by Pfizer ASPIRE Grant WI215080, NIH Award P30 AR070155, and the Rheumatology Research Foundation. JA was supported by Rosalind Russell Medical Research Foundation Bechtel Award, the Arthritis National Research Foundation Award, UCSF Institute for the Rheumatic Diseases, UCSF Research Evaluation & Allocation Committee funded by Esther Memorial fund, and K08 AR072144. We would like to thank Dr. Boris Oskotsky for technical support and the members of the Sirota Lab for useful discussion. We would also like to thank Cindy Lin for her initial work aggregating gene expression datasets and carrying out preliminary analysis of a subset of data.

## Tables

**Table 1.**
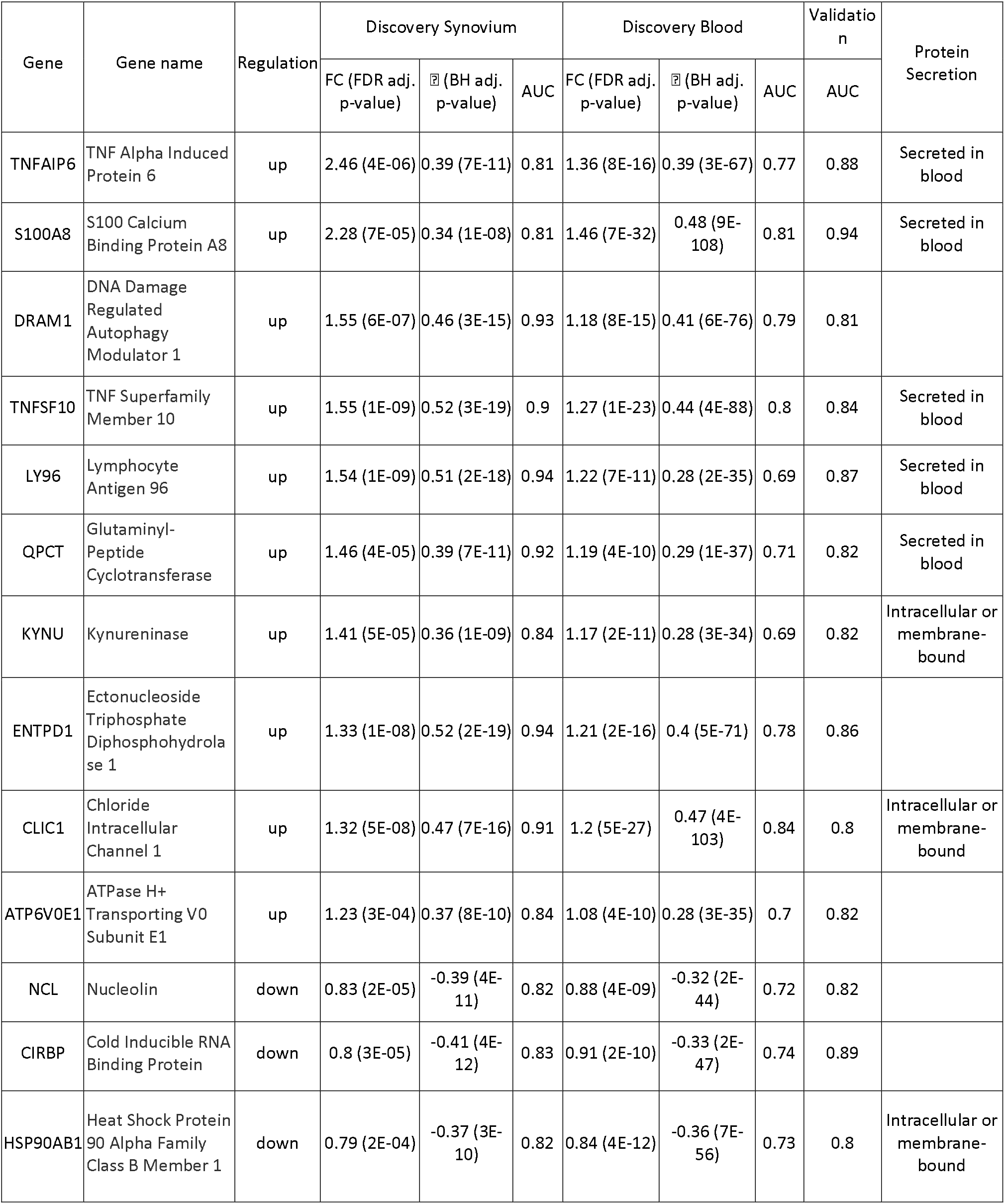
Summary of 13 validated RA Score Panel genes.

## Supplementary Tables

**Supplementary Table 1. Overview of the Discovery and Validation Studies**

**Supplementary Table 2. Treatment Categories**

**Supplementary Table 3. Significantly Differentially Expressed Genes - Synovium**

**Supplementary Table 4. Significantly Differentially Expressed Genes - Blood**

**Supplementary Table 5. Significant Pathways - Synovium**

**Supplementary Table 6. Significant Pathways - Blood**

**Supplementary Table 7. 33 Common DE Genes**

**Supplementary Table 8. 53 Feature Selected Genes**

## Supplementary Figure Legends

**Supplementary Figure 1. Discovery Data Batch Correction, QC and PCA plots**

For synovium and whole blood: a) and b) before batch correction; c) and d) after normalization colored by batch; and f) after normalization colored by treatment type; g) and h) after normalization colored by phenotype

**Supplementary Figure 2. Blood Differential Expression and Pathway Analysis**

DGE analysis in synovial tissue. a) Heatmap and b) PCA plot with DE genes. Reactome pathways for c), d) up- and e), f) down-regulated genes.

**Supplementary Figure 3. Synovium Differential Expression and Pathway Analysis**

DGE analysis in whole blood. a) Heatmap and b) PCA plot with DE genes. Reactome pathways for c), d) up- and e), f) down-regulated genes.

**Supplementary Figure 4. Machine Learning Model Comparison**

AUROC plots for common and feature selected genes. Three models, a Logistic Regression, Elastic Net and Random Forest, were trained on the discovery whole blood data using either common genes or feature selected genes and validated on 5 validation datasets. The summary curves are the averaged curves with bars of standard errors and colored by red. The dashed and solid lines represent synovium and blood data, respectively.

Supplementary Figure 5. Validation Data Batch Correction, QC and PCA Plots

Heatmap and PCA plots of 13 best performing genes on the independent validation a) synovium RNA-seq GSE89408, b) synovium microarray GSE1919, c) whole blood microarray GSE90081, d) PBMC RNA-seq GSE17755 and e) PBMC microarray GSE15573 datasets.

**Supplementary Figure 6. Correlation forest plots with DAS28 for all 13 feature selected genes**

**Supplementary Figure 7. Correlation of DAS score with RA Score for synovium GSE45867 and blood GSE15258, GSE58795, GSE93272 datasets.**

**Supplementary Figure 8. The second set of western blot gels.**

