## Supplementary Figures for "Cross-Tissue Transcriptomic Analysis Leveraging Machine Learning Approaches Identifies New Biomarkers for Rheumatoid Arthritis"

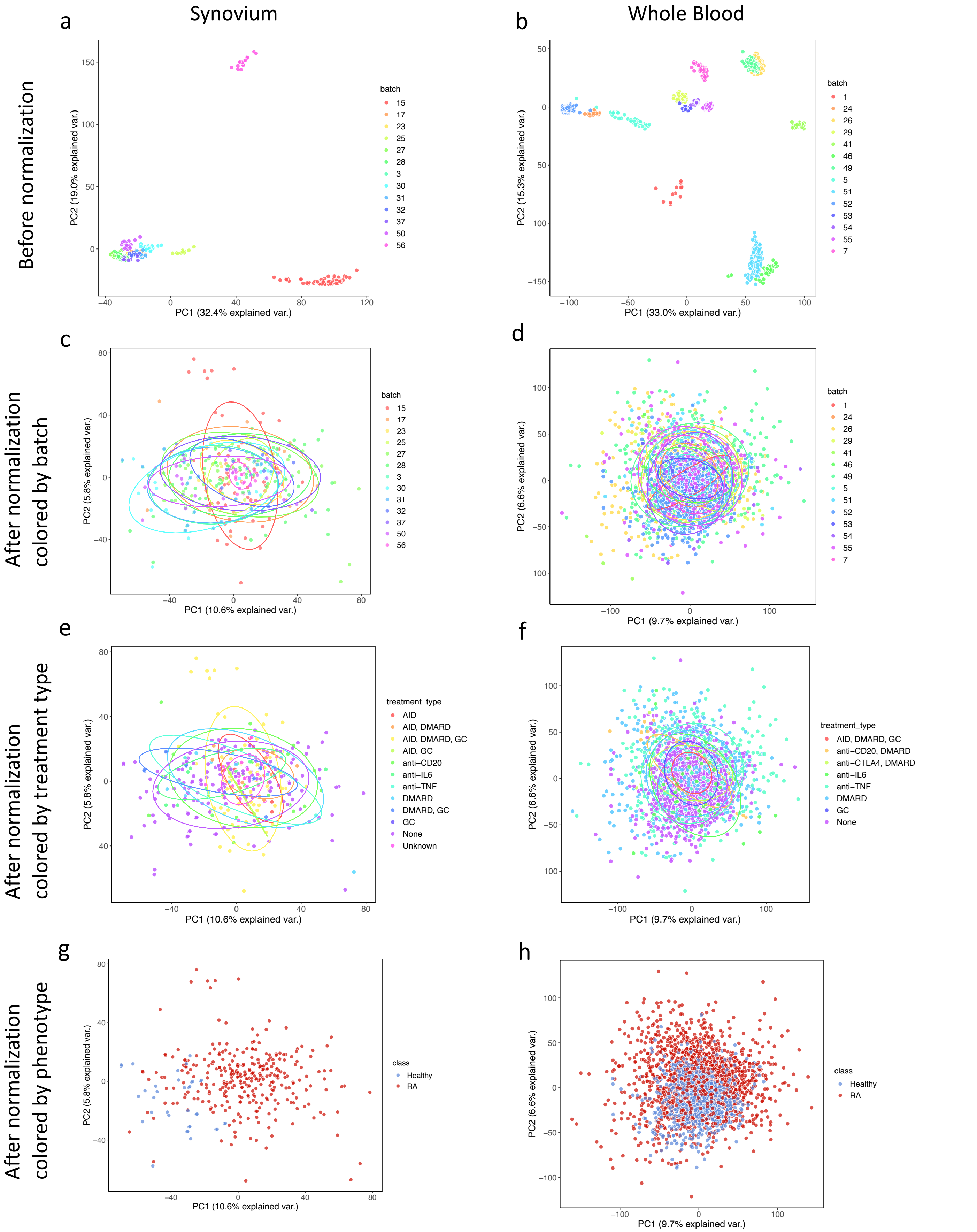


**Supplementary Figure 1. Discovery Data Batch Correction, QC and PCA plots**

For synovium and whole blood: a) and b) before batch correction; c) and d) after normalization colored by batch; e) and f) after normalization colored by treatment type; g) and h) after normalization colored by phenotype.

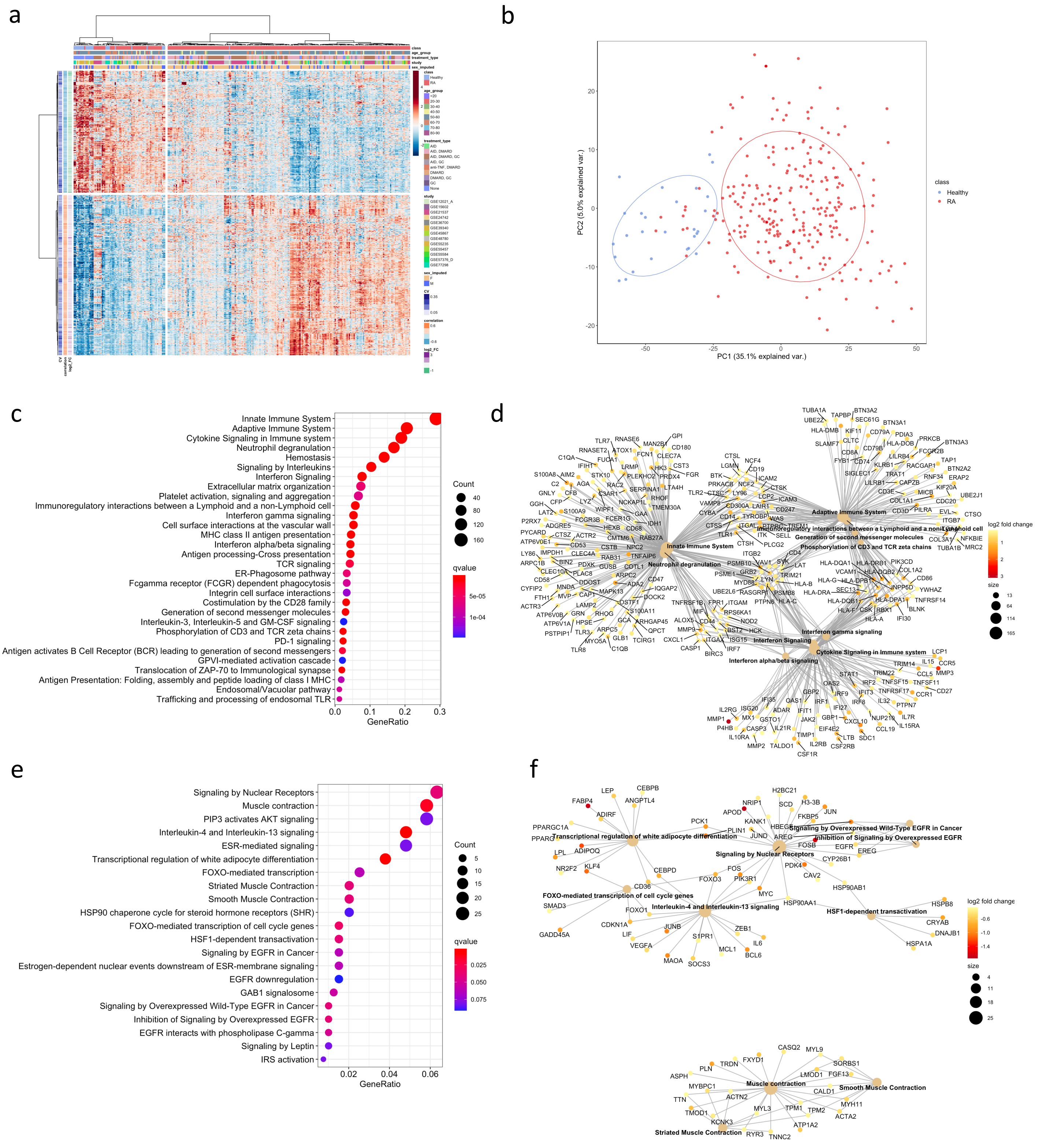


**Supplementary Figure 2. Synovium Differential Expression and Pathway Analysis**

DGE analysis in synovial tissue. a) Heatmap and b) PCA plot with DE genes. Reactome pathways for c) , d) up- and e), f) down-regulated genes.

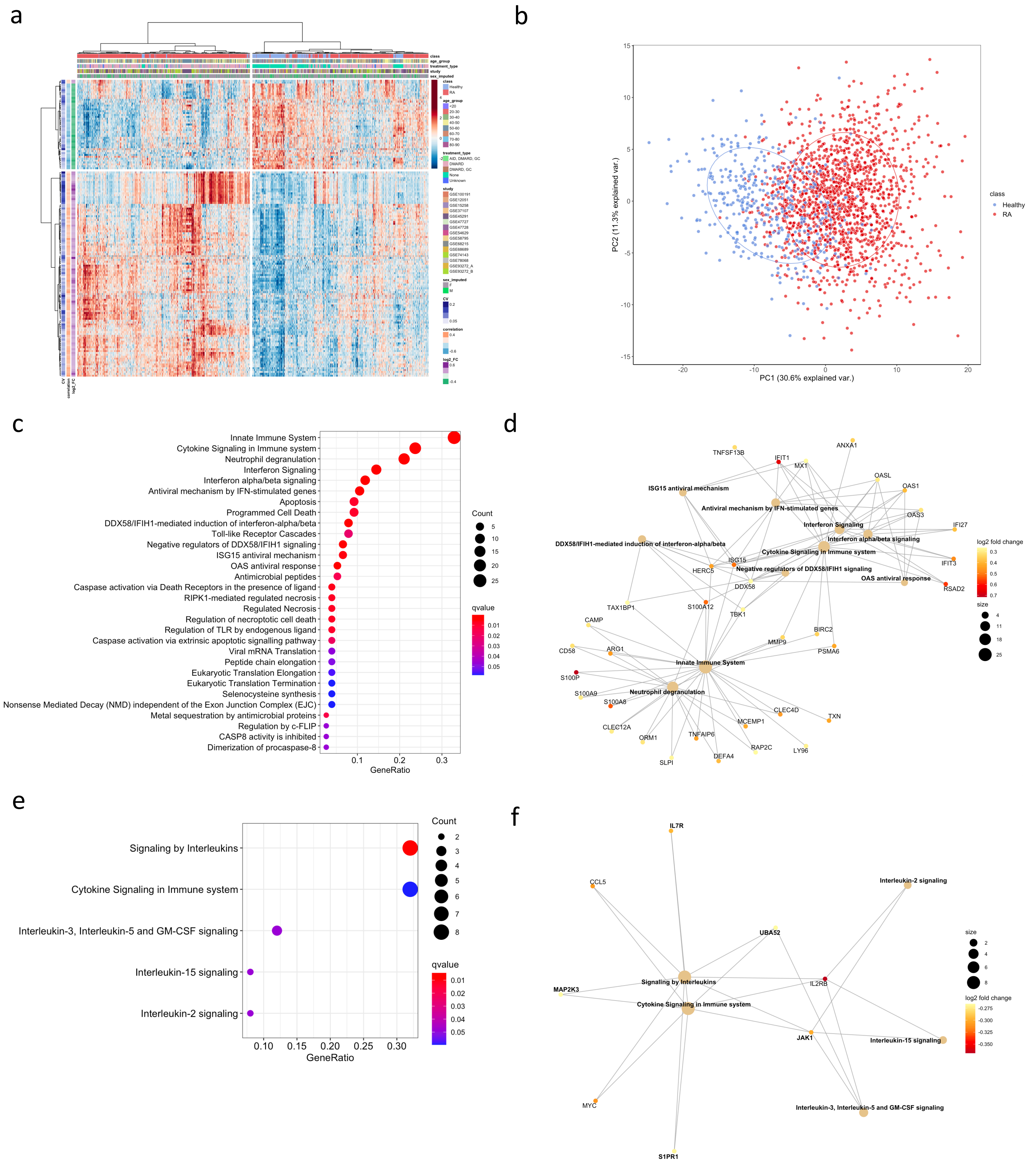


**Supplementary Figure 3. Synovium Differential Expression and Pathway Analysis**

DGE analysis in whole blood. a) Heatmap and b) PCA plot with DE genes. Reactome pathways for c) , d) up- and e), f) down-regulated genes.

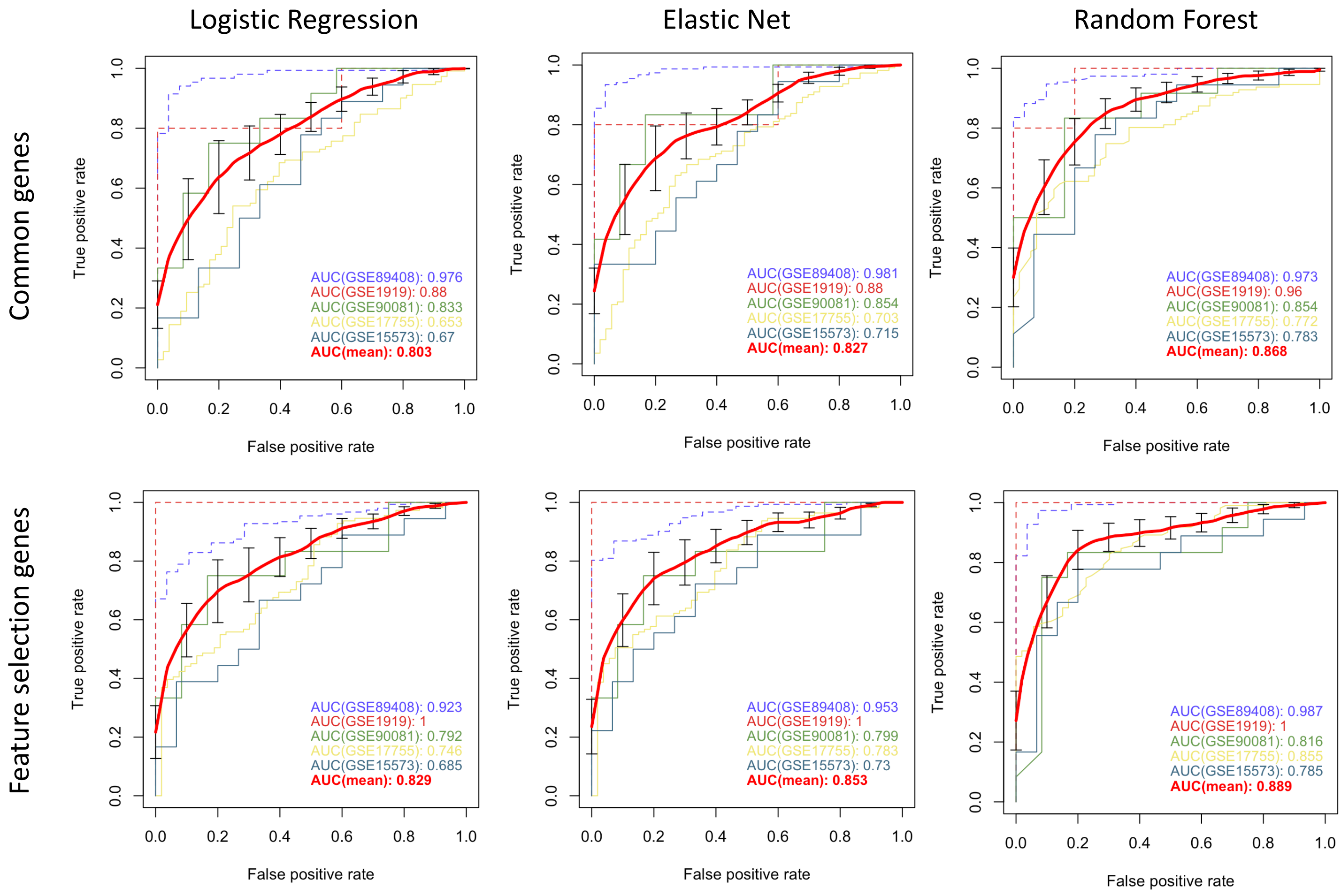


**Supplementary Figure 4. Machine Learning Model Comparison**
AUROC plots for common and feature selected genes. Three models, a Logistic Regression, Elastic Net and Random Forest, were trained on the discovery whole blood data using either common genes or feature selected genes and validated on 5 validation datasets. The summary curves are the averaged curves with bars of standard errors and colored by red. The dashed and solid lines represent synovium and blood data, respectively.

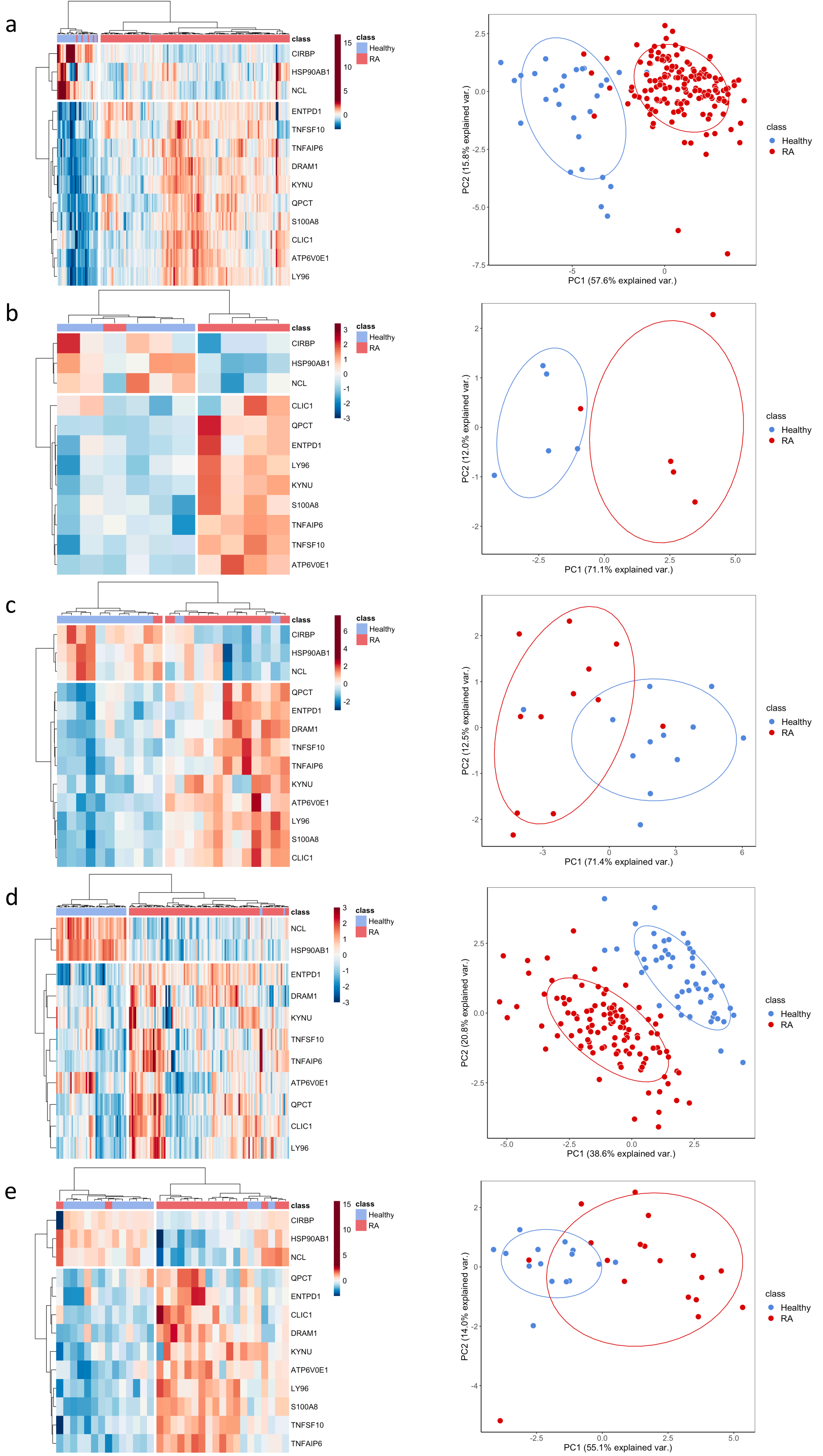


**Supplementary Figure 5. Validation Data Batch Correction, QC and PCA Plots**

Heatmap and PCA plots of 13 best performing genes on the independent validation a) synovium RNA-seq GSE89408, b) synovium microarray GSE1919, c) whole blood microarray GSE90081, d) PBMC RNA-seq GSE17755 and e) PBMC microarray GSE15573 datasets.

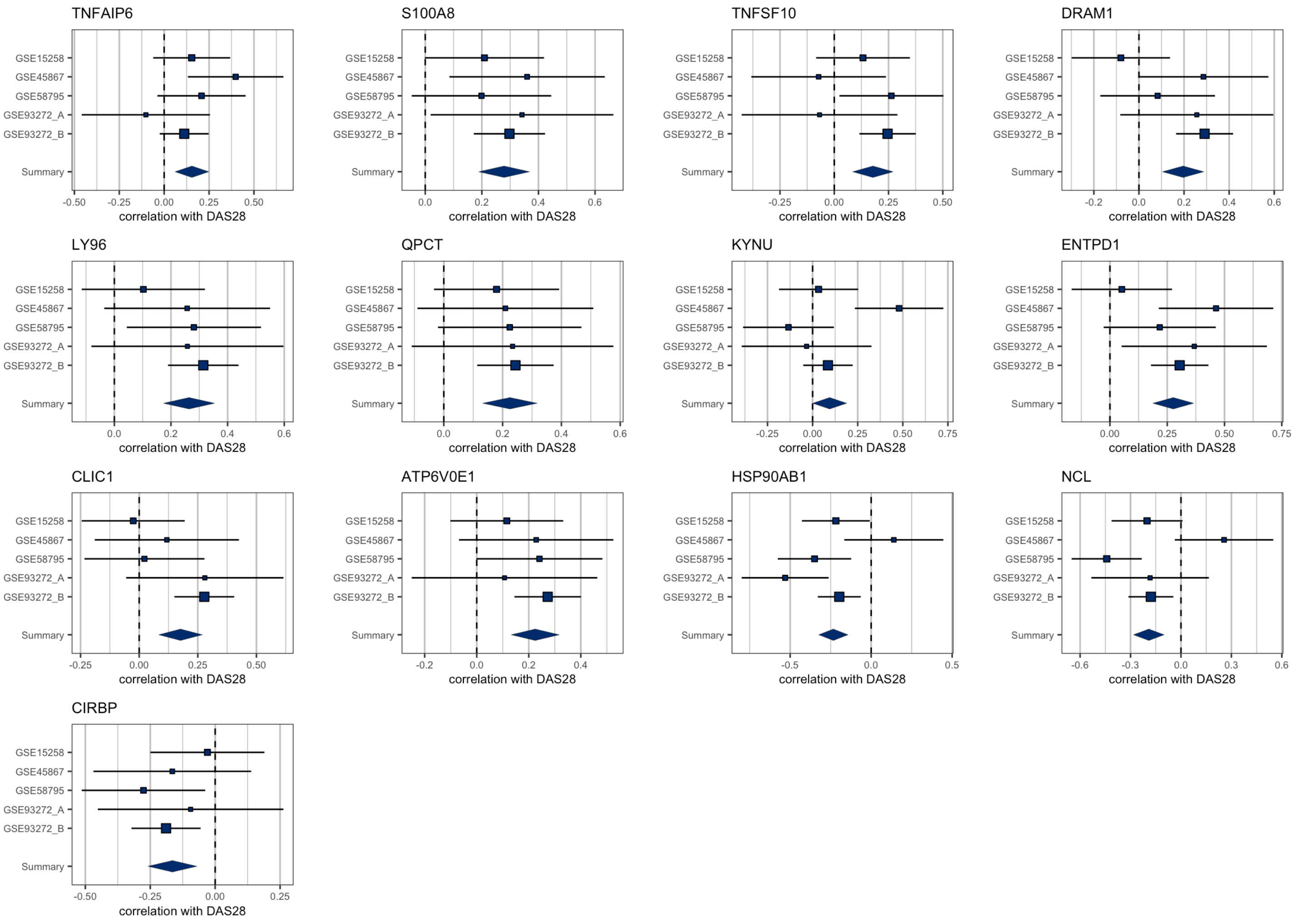


**Supplementary Figure 6. Correlation forest plots with DAS28 for all 13 feature selected genes.**

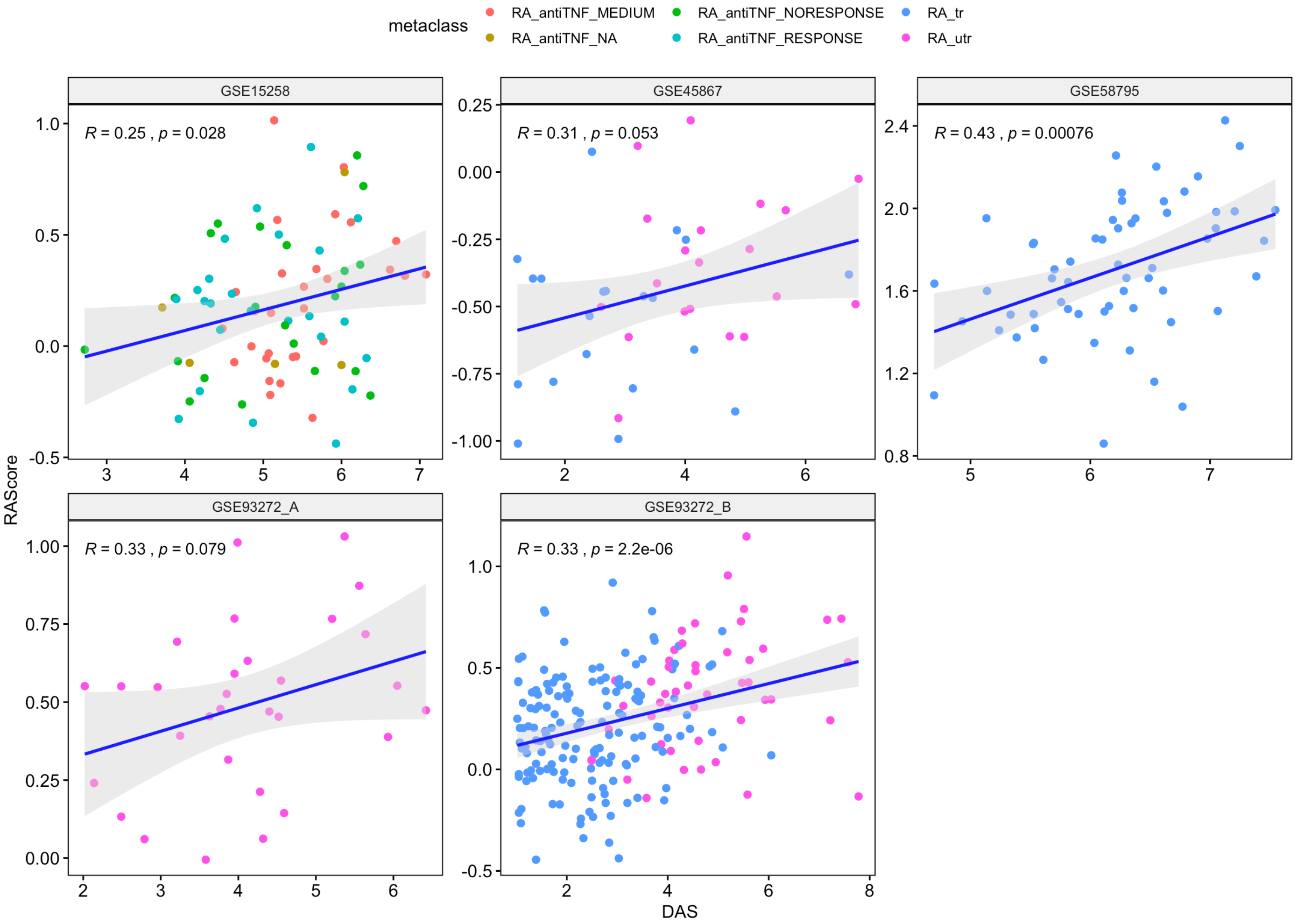


**Supplementary Figure 7. Correlation of DAS score with RA Score for synovium GSE45867 and blood GSE15258, GSE58795, GSE93272 datasets.**

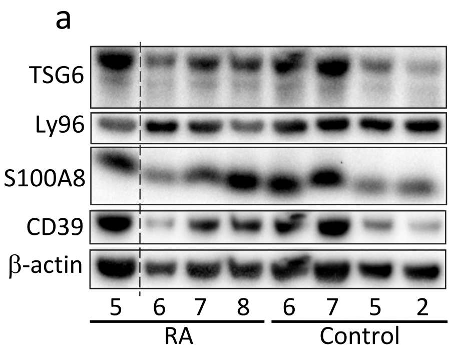


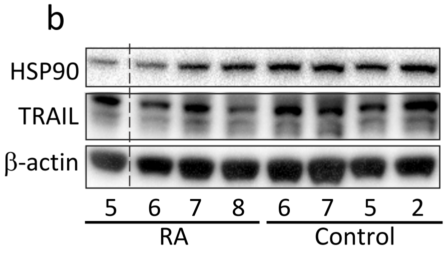


**Supplementary Figure 8. The second set of western blot gels.**
